# MRI-Based Quantification of Central Nervous System Tissue and Cerebrospinal Fluid Volumes in Developing Pigs

**DOI:** 10.1101/2025.10.21.683566

**Authors:** Luke S. Myers, John F. Griffin, Sarah G. Christian, Scott V. Dindot

## Abstract

There is a growing need for alternative animal models to test brain-targeted therapies, and pigs are emerging as a promising option. Their utility, however, depends on reliable estimations of central nervous system (CNS) tissue and cerebrospinal fluid (CSF) volumes, which are essential for translating therapeutic doses between studies in animals and humans. To address this need, we conducted a cross-sectional study of 12 commercial pigs (*Sus scrofa*) across four age groups (2, 5, 11, and 19 weeks). High-resolution magnetic resonance images (MRI) of the brain and spinal cord were acquired using T2-weighted turbo spin echo with fat saturation and short *tau* inversion recovery scans. CNS tissue and CSF volumes were segmented and quantified using 3D Slicer, along with additional anatomical measurements. Our findings reveal notable age-related changes, including spinal CSF volume surpassing brain CSF volume in older pigs, highlighting shifts in CSF distribution that may influence dosing and delivery strategies for CNS-targeted therapies. This study provides a reference for future research using pig models in CNS disease studies and underscores the importance of incorporating brain and spinal CSF and CNS volume data into preclinical models.

## Introduction

Preclinical research on central nervous system (CNS) therapies primarily relies on rodent and non-human primate (NHP) models [1-4]; however, both models have notable limitations. Rodents, while readily accessible, often exhibit poor translatability to humans due to their small size, substantial anatomical differences, and limited predictive value for pharmacokinetic (PK), pharmacodynamic (PD), and toxicological outcomes in humans [5-7]. NHPs are anatomically and physiologically more similar to humans, but their use is increasingly constrained by ethical concerns, high costs, limited availability, and the lack of genetically modified disease models [8].

These challenges have created a growing need for alternative preclinical models, particularly large animal models. Pigs (*Sus scrofa*) are an attractive option due to their anatomical and physiological similarities to humans, cost-effectiveness, and ease of maintenance [9]. They can produce large numbers of offspring in a relatively short period and there is an expanding repertoire of genetically engineered models for CNS disorders, including Huntington’s disease, Batten disease, and Angelman syndrome [7,9-11]. However, to effectively leverage pigs in CNS research, a deeper understanding of their CNS tissue and cerebrospinal fluid (CSF) volumes is required.

Accurate CNS and CSF volume estimates are important for designing preclinical PK/PD and toxicology studies, optimizing translational research, and determining human-equivalent dosing during the preclinical-to-clinical transition [5]. This is especially relevant for CNS-targeted therapies, such as gene therapies, antisense oligonucleotides, and biologics, which are commonly delivered via intracerebroventricular, cisterna magna, or lumbar intrathecal routes [5,6,12]. The translation of doses between preclinical studies in animals and clinical studies in humans primarily involves a scaling model of CSF volume between the animal model and humans. For instance, the human equivalent dose of brain-targeted therapies is 10X of that tested in cynomolgus macaques, as humans and cynomolgus macaques are estimated to have on average 150 ml [13-17] and 15 ml [5] of CSF, respectively.

Despite their utility, current dosing models typically focus solely on brain CSF volume using single average estimates, neglecting the spinal cord’s contribution and developmental changes in CNS and CSF volumes [13-16]. These limitations hinder therapeutic development for neurodevelopmental disorders, where age-related changes in CSF volume—particularly the disproportionate expansion of spinal CSF compared to brain CSF during development—affect therapeutic dosing [18].

The combined CSF and CNS tissue volumes in pigs have never been studied. To address these gaps, we conducted magnetic resonance imaging (MRI) using T2-weighted turbo spin echo (TSE) with fat saturation and short *tau* inversion recovery (STIR) scans on the brains and spinal cords of 12 commercial pigs, ranging in age from 2 weeks (3.7 kg) to 19 weeks (57 kg). We assessed CSF and CNS tissue volumes while obtaining additional anatomical CNS measurements. Our analysis revealed that spinal CSF volume surpasses brain CSF volume in older pigs, emphasizing the importance of accounting for developmental changes. These findings establish a foundational reference for using pig models in CNS disorder studies and underscore the necessity of incorporating both brain and spinal CSF and CNS tissue volume data into preclinical models.

## Methods

### Pig health and housing

This study was approved by the Institutional Animal Care and Use Committee (IACUC) at Texas A&M University. Twelve Yorkshire-Landrace pigs (5 females, 7 males) from 10 litters were included to ensure genetic diversity. The age groups and corresponding ranges in days were as follows: 2 weeks (14 days), 5 weeks (30–37 days), 11 weeks (73–80 days), and 19 weeks (134– 137 days) (**Supplementary Table 1**). Pigs were housed in climate-controlled rooms (74 ± 1°F, 12-hour light/dark cycles, 40–60% humidity). Feed was provided ad libitum until 40 days of age and was subsequently adjusted to 4% and 2% of body weight daily for pigs aged 40–120 days and >120 days, respectively. Animals were group-housed, received no medical treatments, and were confirmed healthy via veterinary assessments prior to scanning.

### Magnetic resonance scanning acquisition

Throughout the MRI scans, pigs were maintained under anesthesia using approximately 3% isoflurane (ASPEN Veterinary Resources) and mechanically ventilated with a Vetland Landmark VSA-2100 Anesthesia System. To minimize motion artifacts, animals were positioned in the supine position. Body temperature was maintained using heating mats and blankets, while physiological parameters, including blood pressure, oxygen saturation, carbon dioxide levels, and body temperature, were continuously monitored.

### Magnetic resonance imaging protocol

All MRI scans were performed using a 3T scanner (Magnetom Verio, Siemens Healthineers, Erlangen, Germany) to obtain high-resolution images of the brain and spinal cord. Two sequences were acquired for each animal: a T2-weighted turbo spin echo (TSE) with fat saturation and short *tau* inversion recovery (STIR). The TSE sequence was acquired in the sagittal plane using the adaptive inline-composed feature, generating a single comprehensive image that included the brain and the entire spinal column. The STIR sequence, obtained in the transverse plane, was optimized for suppressing fat signals, thereby enhancing the visualization of CSF and other fluids. Both sequences leveraged the adaptive inline-composed feature to maintain image consistency across and complete anatomic coverage of the full spinal length.

Scans were conducted over 1.5 to 2 hours, depending on the number of steps required to compose the entire spinal column. Pigs were positioned supine within the MRI scanner, with the brain aligned in the neck coil and additional body matrix and spine coils employed to maximize signal-to-noise ratio across the imaging field. Throughout the procedure, RF pulse settings were standardized to Low SAR for both sequences to ensure safety and image uniformity.

### Sequence parameters

For the TSE sagittal sequence, imaging parameters included a repetition time (TR) of 4800 ms, an echo time (TE) of 87 ms, a field of view (FoV) of 225 mm with 100% phase coverage, a matrix size of 320 mm × 80 mm, and a slice thickness of 2 mm. The flip angle was set to 120 degrees with two averages. The STIR transverse sequence was acquired with a TR of 8830 ms and a TE of 32 ms, an FoV of 125 mm with 100% phase coverage, a matrix size of 256 mm × 100 mm, and a slice thickness of 3 mm. The flip angle was 120 degrees, with parallel imaging (GRAPPA) enabled and an acceleration factor of 2.

### CSF and tissue segmentation

Segmentation and measurements were performed using 3D Slicer software [19], employing the paint and auto-fill tools to segment ventricles and the subarachnoid space in STIR transverse images. For Pig ID 10, T2-weighted sagittal images were used due to STIR image errors. Image contrast and intensity were adjusted to delineate the boundaries of ventricles and subarachnoid space, with surrounding brain and spinal tissues also segmented. Brain CSF was defined as including the lateral, third, and fourth ventricles, as well as the subarachnoid space. Spinal CSF encompassed the subarachnoid space from the foramen magnum to the lumbar cistern. Combined CSF, and combined tissue volumes refer to the sum of the respective brain and spinal volumes, with the foramen magnum serving as the boundary between the brain and spine compartments. All scans were reviewed twice to ensure accuracy and consistency. Subsequent data were graphically visualized using GraphPad Prism (2024).

### CNS measurements

CNS dimensions were assessed using 3D Slicer line and curve tools [19]. Measurements included brain cavity length, height, and width; spinal cord and canal lengths; and cord and cavity heights at key vertebrae. Lengths and heights were measured on T2-weighted sagittal images with fat saturation. Spinal cord length was measured from the foramen magnum to the caudal point of the conus medullaris, which is long and tapered in pigs and ends at approximately S2 [20]. Spinal canal length was measured from the foramen magnum to the caudal aspect of the lumbar cistern. Brain cavity length was measured as the maximal distance from the cribriform plate to the occipital bone, while brain cavity height was measured orthogonally at the level of the sella turcica. Brain cavity width was determined in STIR transverse plane images at the widest point in the slice containing the rostral colliculi to ensure consistency across animals. Spinal cord height was measured at C1, T1, and L1, while spinal canal height was recorded at the widest point of each vertebra. Spinal subarachnoid space height was calculated by subtracting the spinal cord height from the spinal canal height and dividing the result by two. Additionally, the length of the CNS was measured from the center of the eye to the first palpable sacral vertebra in MRI images, excluding pig ID 11 due to image limitations. This metric was preferred over snout-to-tail length due to variability in snout length and tail docking practices.

### Vertebrae count

Vertebrae were manually counted from C1 to C7, followed by thoracolumbar vertebrae to determine the total thoracic and lumbar count. However, individual thoracic and lumbar vertebrae counts were not feasible due to limited scan coverage focused on the brain and spinal cord, which obscured rib visibility and the thoracic-lumbar boundary.

### Comparison of 19-week-old pig CNS measurements against other species

Publicly available data from healthy humans, macaques, rats, and mice were included, using only averages reported or calculated from raw data to avoid extrapolation errors. For studies reporting both sexes, data were averaged, and regional measurements were combined as needed. Only adult specimens were included, and for studies with multiple ages, closely aligned ages were averaged. All lengths were standardized to centimeters (cm) and volumes to cubic centimeters (cm^3^). Brain tissue volume was derived by subtracting CSF volume from intracranial volume when both were reported.

## Results

### Pig CNS measurements across multiple age groups

To evaluate CSF volume, CNS tissue volume, and other anatomical CNS measurements, MRI scans were performed on 12 Yorkshire/Landrace pigs across four age groups: 2 weeks (n = 3), 5 weeks (n = 3), 11 weeks (n = 3), and 19 weeks (n = 3) (**Table 1**). These age groups represent brain development ranging from 42.3% at 2 weeks, 50–53.9% at 5 weeks, 71.9–76% at 11 weeks, and 93% at 19 weeks of maximum brain growth [21]. All animals tolerated the MRI protocol well, producing high-quality images suitable for analysis without significant artifacts (e.g., flow, motion, aliasing). Image segmentation and volume measurements were performed using 3D Slicer [19] (**Figure 1 A-F**). Brain CSF measurements included the lateral, third, and fourth ventricles as well as the subarachnoid space. Spinal CSF encompassed the subarachnoid space from the foramen magnum to the lumbar cistern. The foramen magnum was used as the boundary between brain and spinal compartments, and combined CSF or combined CNS tissue volume represented the sum of brain and spine volumes.

**Table 1.** Study groups.

| **Pig ID** | **Age Group (Weeks)** | **Sex** | **Weight (kg)** | **CNS Length (cm)** |
| --- | --- | --- | --- | --- |
| 1 | 2 | M | 3.7 | 37.4 |
| 2 | 2 | F | 3.8 | 37.2 |
| 3 | 2 | M | 4.3 | 38.3 |
| 4 | 5 | F | 7.8 | 46.4 |
| 5 | 5 | M | 7.2 | 47.2 |
| 6 | 5 | F | 7.3 | 48.1 |
| 7 | 11 | M | 26.0 | 68.8 |
| 8 | 11 | M | 33.0 | 74.1 |
| 9 | 11 | F | 27.0 | 72.5 |
| 10 | 19 | F | 57.0 | 86.3 |
| 11 | 19 | M | 47.0 | N/A |
| 12 | 19 | M | 56.0 | 87.7 |

M = male, F = female, CNS = central nervous system, and N/A = data not available.

**Figure 1.**
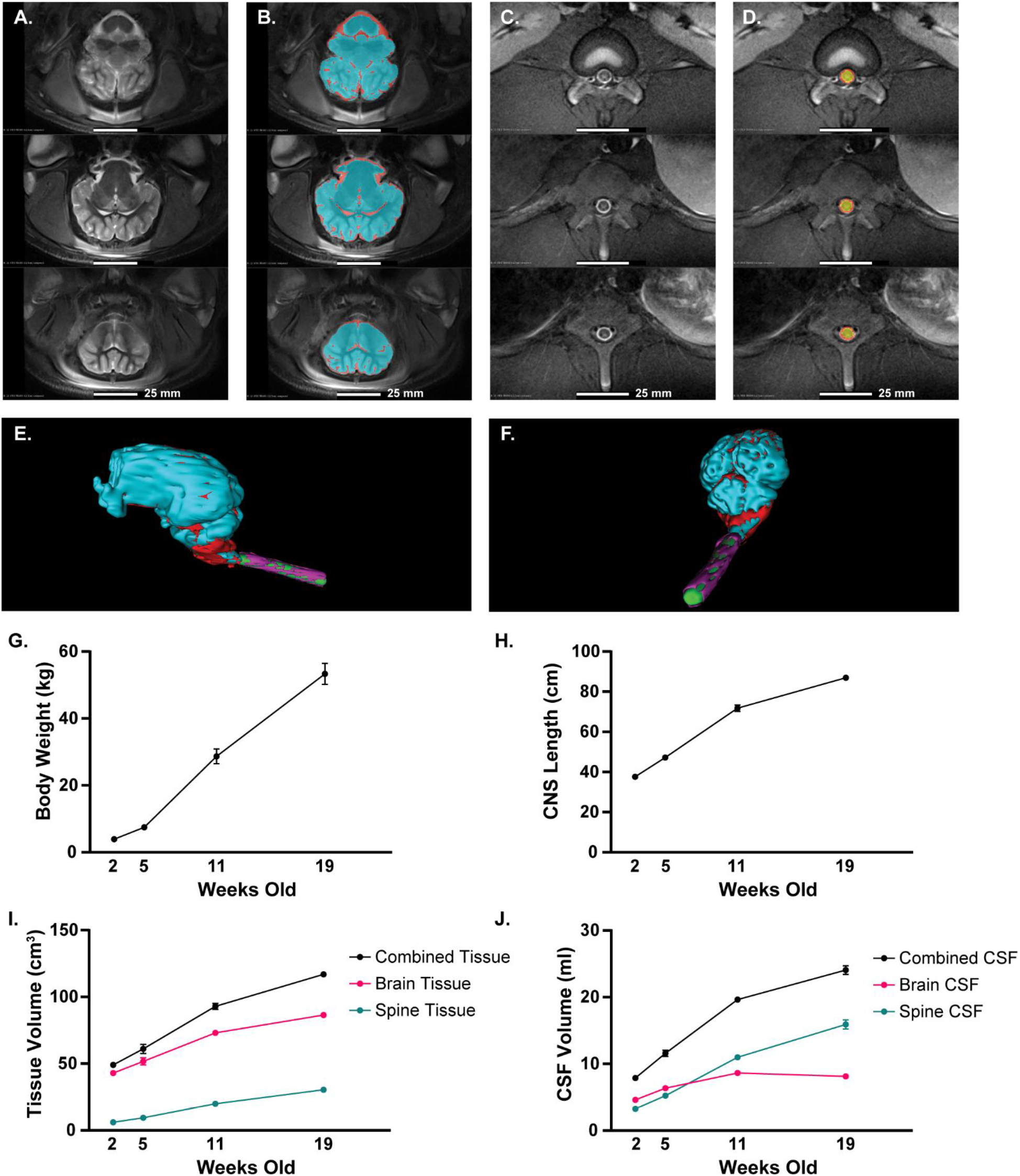
Segmentation of CSF and CNS tissue, and developmental changes in pigs. Representative MRI slices and 3D renderings illustrate CSF and CNS tissue segmentation in the brain and spinal cord of an 11-week-old pig (ID 8). (**A, B**) Transverse STIR MRI slices of the brain showing original images (**A**) and segmented CSF (red) and CNS tissue (blue) (**B**). (**C, D**) Transverse STIR MRI slices of the spinal cord displaying original images (**C**) and segmented CSF (red) and CNS tissue (yellow) (**D**). (**E, F**) 3D renderings of segmented CSF (red/purple) and CNS tissue (blue/green). (**G**) Mean body weight across ages (2, 5, 11, and 19 weeks). (**H**) Mean CNS length across ages. (**I**) Mean CNS tissue volume across ages. (**J**) Mean CSF volume across ages. Segmentations were performed using 3D Slicer software [19]. Data presented as mean ± SEM and visualized using GraphPad Prism (2024). Abbreviations: cerebrospinal fluid (CSF), and central nervous system (CNS).

The largest increase observed between the 2-week-old and 19–week-old groups was in body weight, with an approximately 13-fold increase from an average of 3.9 kg to 53.3 kg (**Figure 1G**). In contrast, increases in CNS length, combined CSF volume, and combined tissue volume were more moderate over the same period. CNS length showed an approximately 2.3-fold increase, from 37.6 cm at 2 weeks to 87 cm at 19 weeks (**Figure 1H**). This was closely mirrored by the combined CNS tissue volume, which increased approximately 2.4-fold, from 49 cm^3^ to 117 cm^3^ (**Figure 1I**). Combined CSF volume exhibited a slightly greater increase, with a 3-fold rise from 7.9 ml at 2 weeks to 24.1 ml at 19 weeks (**Figure 1J**). **Figure 2** visually illustrates the increase in CSF volume and CNS tissue between 2 and 19 weeks in pigs, with additional details on individual measurements, including sex, voxel counts, and surface areas, provided in **Supplementary Table 1**.

**Figure 2.**
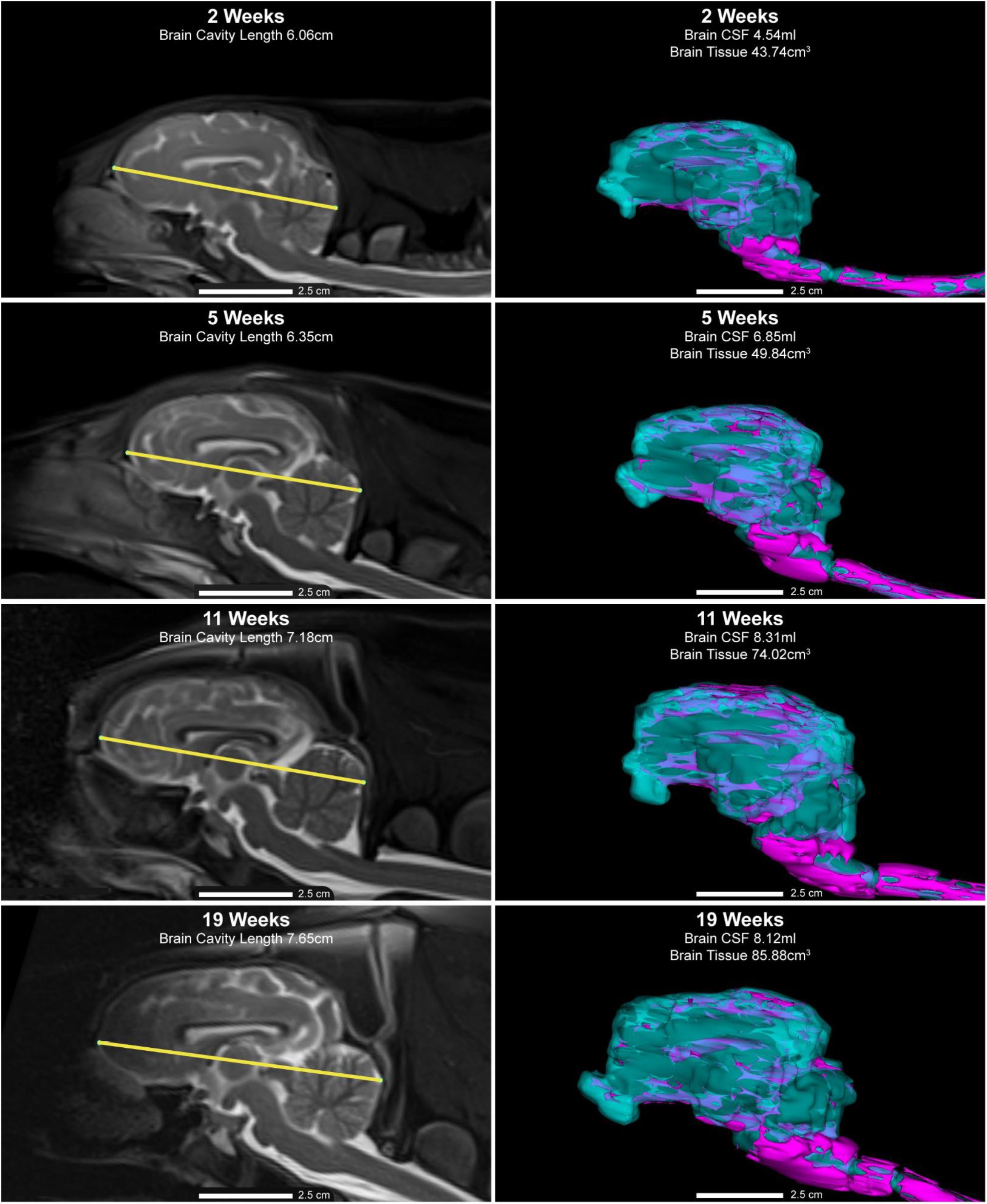
Brain growth and development across ages. Representative MRI scans and 3D renderings from one pig in each age group: Pig ID 2 (2 weeks), Pig ID 5 (5 weeks), Pig ID 9 (11 weeks), and Pig ID 11 (19 weeks). Left panels: T2-weighted sagittal images with brain cavity length (yellow line) measured from the cribriform plate to the occipital bone. Right panels: 3D renderings of brain and spinal tissue (blue) and cerebrospinal fluid (CSF, purple). Measurements and 3D renderings were performed using 3D Slicer [19]. Abbreviations: cerebrospinal fluid (CSF).

Understanding these developmental changes is critical, as they can influence therapeutic dosing strategies and the efficacy of drug delivery at different administration sites. The most notable change was observed in the ratio of spine CSF volume to brain CSF volume. At 2 weeks, spine CSF volume was approximately 70% of brain CSF volume, but by 19 weeks, spine CSF volume had nearly doubled, reaching 195% of brain CSF volume (**Table 2**). Additional comparisons of CNS and CSF volumes at each age are provided in **Table 2**, with the largest changes observed in the spinal compartment.

**Table 2.** Proportional changes in CNS and CSF volumes during development.

| Comparisons of Volumes | 2<br>Weeks | 5<br>Weeks | 11<br>Weeks | 19<br>Weeks |
| --- | --- | --- | --- | --- |
| Spine CSF/Brain CSF (%) | 70.8 | 82.2 | 127.3 | 195.3 |
| Spine Tissue/Brain Tissue (%) | 13.9 | 18.3 | 27.3 | 35.2 |
| Brain CSF/Brain Tissue (%) | 10.5 | 12.3 | 11.8 | 9.4 |
| Spine CSF/Spine Tissue (%) | 53.9 | 55.4 | 55.3 | 52.2 |
| Combined CSF/ Combined CNS Tissue (%) | 16.1 | 19.0 | 21.1 | 20.6 |

CSF = cerebrospinal fluid, and CNS = central nervous system.

### Additional pig anatomical measurements across multiple age groups

Additional anatomical measurements revealed size increases across multiple parameters, although many were not as substantial as those observed for CNS length, CSF volume, and tissue volume (**Table 3**). Similar to these metrics, spinal cord length showed a 2.3-fold increase, from 30.9 cm at 2 weeks to 69.6 cm at 19 weeks (**Figure 3A**). In contrast, brain length and width exhibited more modest 1.3-fold and 1.2-fold changes (**Figure 3B-C**). Brain length increased from 6.1 cm at 2 weeks to 7.8 cm at 19 weeks, while brain width increased from 4.6 cm to 5.6 cm over the same period (**Table 3**). Comparable size increases were observed across other anatomical measurements, with the greatest changes consistently occurring in the spinal region (**Table 3**).

**Table 3.** Average anatomical CNS measurements in pigs.

|  | **2 Weeks** | **5 Weeks** | **11 Weeks** | **19 Weeks** |
| --- | --- | --- | --- | --- |
| **Spinal Cord Length (cm)** | 30.903 | 38.920 | 59.490 | 69.610 |
| **Spinal Canal Length (cm)** | 33.047 | 42.110 | 63.720 | 78.480 |
| **Brain Cavity Length (cm)** | 6.103 | 6.337 | 7.288 | 7.793 |
| **Brain Cavity Width (cm)** | 4.598 | 4.858 | 5.334 | 5.576 |
| **Brain Cavity Height (cm)** | 3.335 | 3.506 | 3.873 | 4.186 |

**Figure 3.**
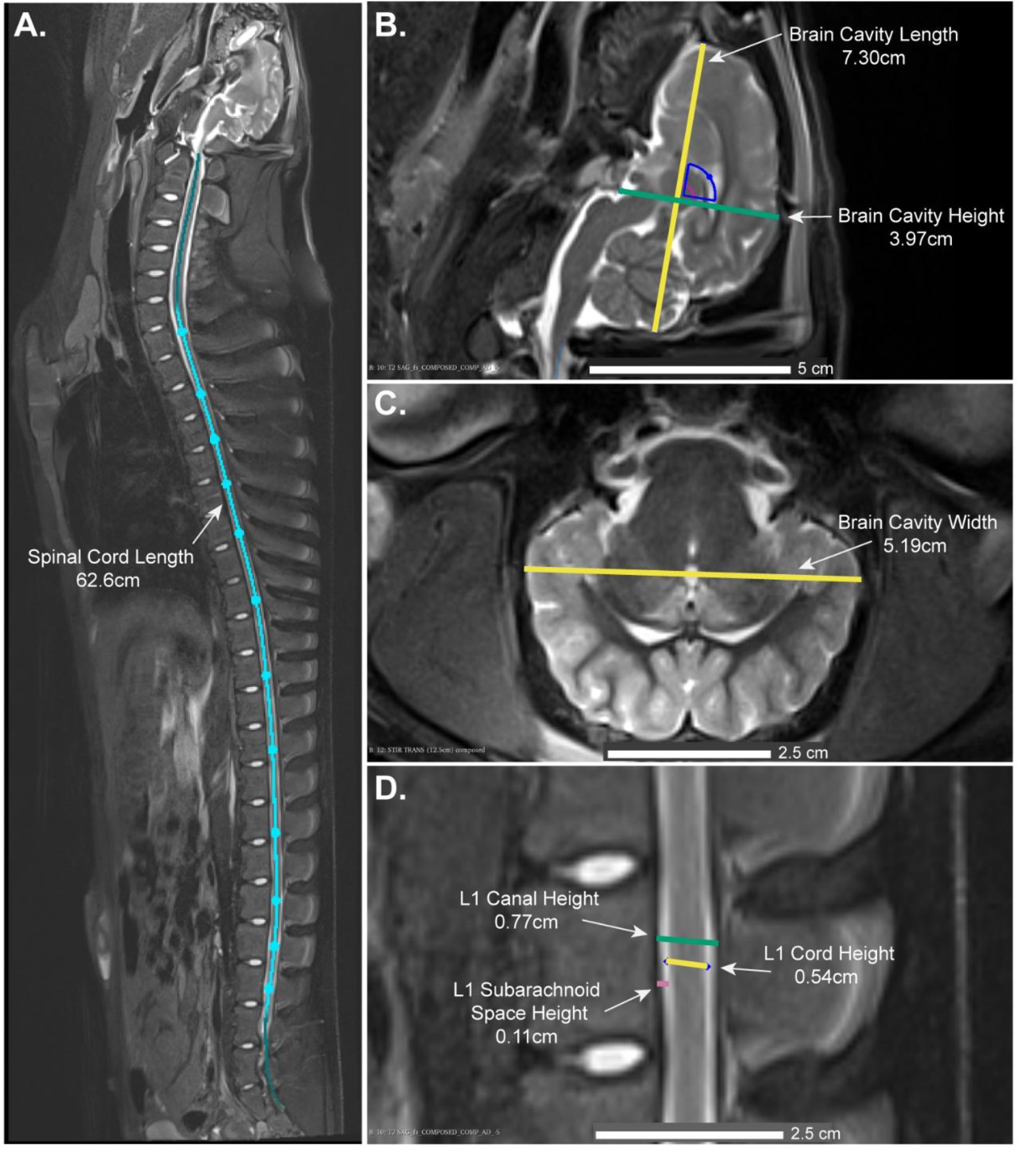
CNS measurements taken in pigs. Representative MRI images illustrating CNS measurement methods in a randomly selected pig (ID 8) from the 11-week-old age group. Panels A, B, and D are T2-weighted sagittal MR images, and panel C is a transverse short tau inversion recovery (STIR) image. (**A**) Spinal cord length measured from the foramen magnum to the caudal end of the conus medullaris (S1–S2). (**B**) Brain cavity dimensions: maximal brain cavity length measured from the cribriform plate to the occipital bone, and brain cavity height measured orthogonally at the level of the sella turcica. (**C**) Brain cavity width measured at the widest point in the slice containing the rostral colliculi. (**D**) L1 spinal cord, spinal canal, and L1 subarachnoid space height measured as the maximal height at the L1 spinal region. Measurements were performed using 3D Slicer [19].

Further details on individual measurements and vertebrae counts are available in **Supplementary Table 1**.

We also obtained measurements for the C1, T1, and L1 vertebrae (**Figure 3D**). The average spinal cord height at C1 showed a 1.5-fold increase, from 0.5 cm at 2 weeks to 0.74 cm at 19 weeks (**Table 4**). In comparison, the average spinal cord height at L1 exhibited a slightly greater 1.8-fold increase, from 0.38 cm to 0.676 cm over the same period (**Table 4**). Interestingly, the subarachnoid space height did not increase proportionally. At C1, the subarachnoid space height only showed a 1.2-fold increase, from 0.12 cm to 0.15 cm, while at L1, it increased 1.4-fold, from 0.1 cm to 0.13 cm (**Table 4**).

**Table 4.** Average C1, T1 and L1 vertebrae measurements.

|  | **2 Weeks** | **5 Weeks** | **11 Weeks** | **19 Weeks** |
| --- | --- | --- | --- | --- |
| **C1 Spinal Cord Height (cm)** | 0.496 | 0.577 | 0.623 | 0.740 |
| **T1 Spinal Cord Height (cm)** | 0.491 | 0.574 | 0.677 | 0.723 |
| **L1 Spinal Cord Height (cm)** | 0.376 | 0.466 | 0.540 | 0.676 |
| **C1 Canal Height (cm)** | 0.739 | 0.855 | 0.913 | 1.029 |
| **T1 Canal Height (cm)** | 0.704 | 0.881 | 0.973 | 1.043 |
| **L1 Canal Height (cm)** | 0.567 | 0.655 | 0.785 | 0.935 |
| **C1 Subarachnoid Space Height (cm)** | 0.121 | 0.139 | 0.145 | 0.145 |
| **T1 Subarachnoid Space Height (cm)** | 0.107 | 0.154 | 0.148 | 0.160 |
| **L1 Subarachnoid Space Height (cm)** | 0.095 | 0.095 | 0.122 | 0.129 |

Further details on individual measurements are available in **Supplementary Table 1**.

### Weight and CNS length correlations for CNS measurement estimations

We investigated whether body weight and CNS length (distance from the eye to the first palpable sacral vertebra) could predict other CNS measurements, such as CSF volume and CNS tissue volume, to facilitate the application of these data in future studies. Correlations were analyzed between CNS measurements and both body weight and CNS length across all age groups (**Supplementary Table 2**). Both metrics demonstrated strong predictive value; however, CNS length consistently showed stronger correlations. These findings suggest that CNS length may provide more reliable estimations, particularly in scenarios where body weight is affected by diet or growth variability.

## Discussion

This study provides new insights into the dynamics of CSF and CNS tissue volumes during pig development, with important implications for dose translation in preclinical studies. Specifically, it reveals age-dependent changes in the relative proportions of CSF in the brain and spine, underscoring the importance of accounting for both compartments when estimating drug dosages based on CSF volume. Notably, spinal CSF volume was lower than brain CSF volume in 2 and 5-week-old pigs but surpassed brain CSF volume by 11 and 19 weeks of age. This transition highlights the limitations of simple scaling models, which often treat the brain as an isolated entity and overlook the interplay between brain and spinal CSF compartments [5,12,16]. Using a single estimated CSF volume across species or age groups may result in disproportionately higher drug exposure in the brain of younger animals compared to older ones. These findings emphasize the need for CNS drug delivery strategies to consider age-specific CSF dynamics, particularly for therapies targeting specific CNS regions.

To validate the translational relevance of pigs as models for studying CNS disorders, we compared CNS metrics of 19-week-old pigs with those of adult humans, macaques, rats, and mice. Data from 53 publicly available studies (**Supplemental Bibliography**) were analyzed, including metrics such as age, CSF volume, CNS tissue volume, and spinal cord length (**Table 5 and Supplementary Table 3**). The brain tissue volume of pigs (86.5 cm^3^) closely approximated that of macaques (88.3 cm^3^), and the combined CSF volume in pigs (24.1 ml) was similarly comparable to macaques (21.4 ml). Interestingly, spinal tissue volume in pigs (30.5 cm^3^) was more similar to humans (20 cm^3^) than to macaques (4.5 cm^3^). However, pigs exhibited a longer spinal cord (69.6 cm) compared to humans (41.4 cm) and macaques (23.2 cm). This is partially explained by their long and tapered conus medullaris ending at approximately the S2 vertebrae, as compared to L1 in humans [20] and L4 in macaques [22]. As expected, rodents (rats and mice) displayed CNS metrics that were markedly different from those of pigs, humans, and macaques (**Table 5**).

**Table 5.**
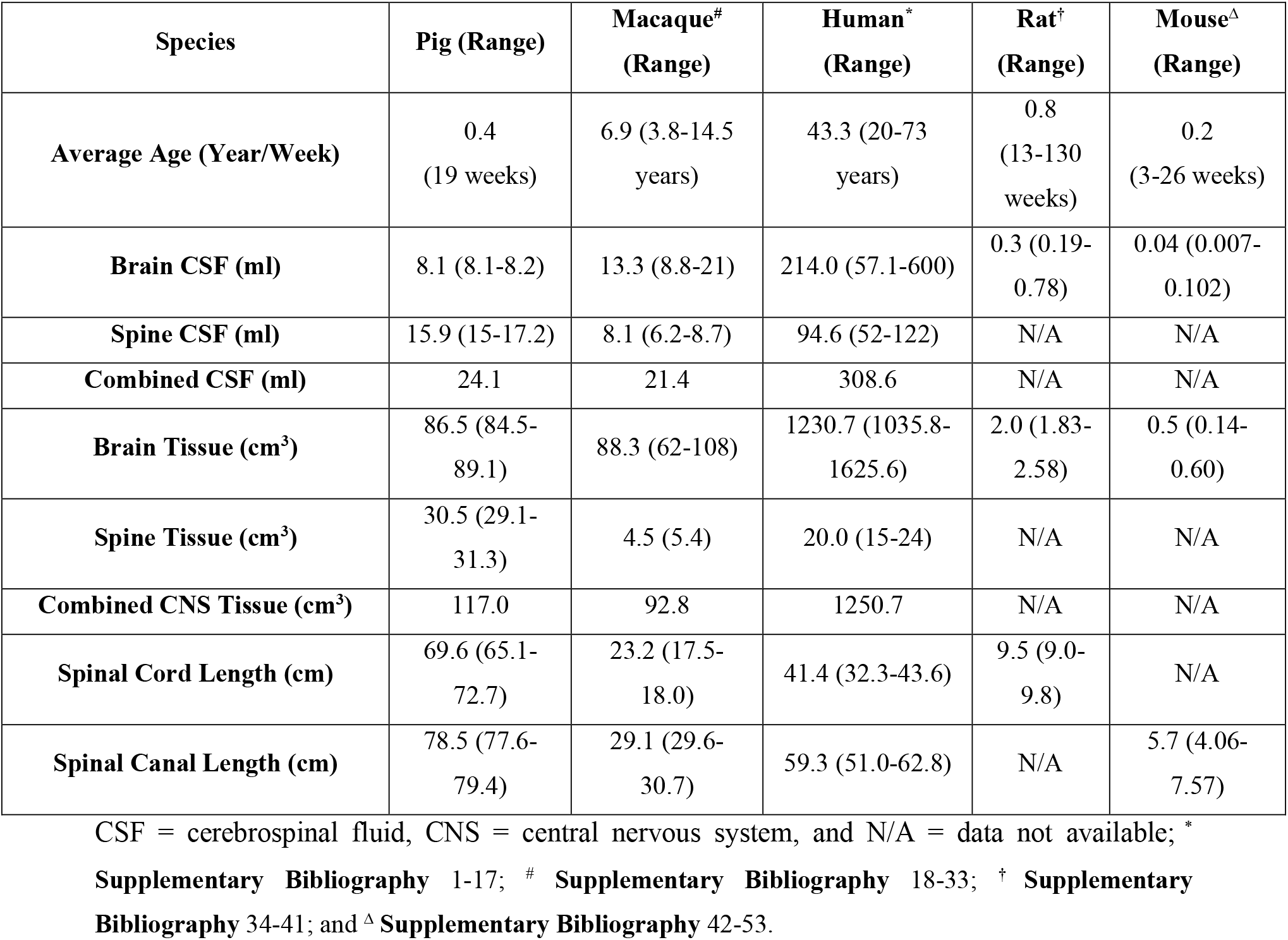
Comparison of multiple CNS measurements in 19-week-old pigs against the averages for humans, macaques, rats, and mice.

The proportional differences in CNS metrics among pigs, macaques, and humans, as summarized in **Table 6**, provide further insight into their translational utility. The ratio of brain CSF volume to brain tissue volume was relatively consistent across pigs (90.6%), macaques (84.9%), and humans (82.6%). Similarly, the ratio of combined CSF volume to total CNS tissue volume was comparable: 79.4% in pigs, 77% in macaques, and 75.3% in humans. However, spine-specific metrics revealed notable differences. The proportion of spine CSF to spine tissue volume was significantly higher in humans (373.4%) than in pigs (47.8%) or macaques (77.6%). Additionally, the proportion of spine CSF to brain CSF was highest in pigs (95.3%), compared to macaques (39.5%) and humans (55.8%) (**Table 6**).

**Table 6.** Mean comparison of relative differences in CNS measurements across pigs, macaques and humans.

| **Comparisons of Volumes** | **Pig** | **Macaque** | **Human** |
| --- | --- | --- | --- |
| **Brain CSF/Brain Tissue (%)** | 90.6 | 84.9 | 82.6 |
| **Spine CSF/Spine Tissue (%)** | 47.8 | 77.6 | 373.4 |
| **Combined CSF/Combined CNS Tissue (%)** | 79.4 | 77.0 | 75.3 |
| **Spine CSF/Brain CSF (%)** | 95.3 | 39.5 | 55.8 |
| **Spine Tissue/ Brain Tissue (%)** | 64.8 | 94.9 | 98.4 |

CSF = cerebrospinal fluid, and CNS = central nervous system

The anatomical differences observed between pigs and macaques should be carefully considered when selecting appropriate models for translational research. Although macaques are often preferred in preclinical research due to their close phylogenetic relationship to humans, our findings suggest that pigs may better replicate human spinal metrics. This makes pigs a particularly suitable model for studies that need to account for both brain and spine dynamics, especially in pediatric research where small juvenile macaques would otherwise be used. Furthermore, pigs offer several logistical and ethical advantages, including lower cost, rapid growth, ease of care, and greater availability, potentially making them a more practical and scalable option for CNS research.

A translational concern previously highlighted in the literature [12,23], and further supported by our findings, is the likely underestimation of CSF volumes in humans and macaques. Standard simple scaling models often estimate CSF volumes as 150 ml for humans [13-17] and 15 ml for macaques [5], typically accounting only for brain CSF. However, our findings indicate that the total CSF volume, including spinal contributions, is approximately 309 ml in humans and 21 ml in macaques. This significant discrepancy underscores the risk of dosing inaccuracies in translational studies when spinal CSF contributions are overlooked.

A key strength of this study is the quantification of both brain and spinal CSF and tissue volumes, along with additional CNS anatomical metrics, within the same animal. To our knowledge, no previous study has collected all these measurements from a single specimen; existing knowledge is typically derived from separate investigations involving different animals and methodologies, limiting the ability to assess relationships among these parameters. Furthermore, this work provides a cross-species comparison of CNS volumetric and anatomical metrics across multiple translationally relevant species, addressing a key gap in the current literature. Notably, despite the extensive use of rodents in biomedical research, we identified surprisingly limited CNS anatomical data for these species, particularly concerning the spinal cord, which may further constrain their translational relevance to human CNS research.

This study is not without limitations. The lack of a reference standard for CSF volume measurements and the resolution constraints of MRI may affect the precision of small-scale structures. Nevertheless, these limitations are unlikely to substantially affect the overall accuracy of our findings. Future studies should aim to refine imaging techniques and include older and larger pigs to better assess age-related changes in CNS metrics. Additionally, breed-specific differences should be explored to further enhance pigs’ utility as translational models.

## Conclusion

This study highlights the dynamic changes in CSF and CNS tissue volumes during pig development, underscoring the importance of considering these shifts when designing CNS-targeted therapies. By comparing CNS metrics across pigs, humans, macaques, and rodents, we demonstrate that pigs serve as a robust and translationally relevant model for studying CNS dynamics. These findings challenge traditional simple scaling models and support the need for revised dosing strategies that account for both brain and spinal CSF contributions in pediatric and adult CNS research. The anatomical and physiological insights provided by this study enhance the translational utility of preclinical models, ultimately advancing the development of more effective CNS therapies.

## Supporting information

Supplementary Bibliography

## Acknowledgments

We thank the Translational Imaging Center at the Texas A&M Institute for Preclinical Studies for providing imaging resources and performing the MRI scans. We are grateful to Sarah Frazier for her assistance with MRI acquisition. We also appreciate the support of the staff from the Comparative Medicine Program and the Texas Institute for Preclinical Studies at Texas A&M.

Additionally, we acknowledge the use of ChatGPT, version 2, developed by OpenAI, for assistance with proofreading, refining language, and improving the clarity and readability of the manuscript text. The authors confirm that no data or scientific content was generated by the language model and take full responsibility for the accuracy and integrity of the final manuscript.

## Author Contributions

Author Contributions: Conceptualization, LSM and SVD; Data curation, LSM and JFG; Formal analysis, LSM and JFG; Funding acquisition, SVD; Investigation, LSM and JFG; Methodology, LSM and JFG; Project administration, LSM and SGC; Resources, JFG; Supervision, SGC and SVD; Validation, JFG and SVD; Visualization, LSM; Writing—original draft, LSM; Writing— review and editing, LSM, JFG, SGC, and SVD.

